# In-Solution Characterization of Biomolecular Interaction Kinetics under Native Conditions

**DOI:** 10.1101/2025.03.27.645748

**Authors:** Philipp Willmer, Emil G. P. Stender, Kritika Sahni Ray, Adam C. Hundahl, Rodolphe Marie, Henrik Jensen

## Abstract

Characterizing the kinetics of biomolecular interactions is fundamental for understanding biological mechanisms, devel-oping novel drugs for advancing healthcare and for optimizing processes in protein engineering. Although modern sur-face-based methods have advanced our understanding of protein-protein and protein-ligand kinetics, they rely on immobilized samples, preventing the study of interactions under native conditions and leading to an incomplete understanding. In this work, we propose a paradigm shift by introducing a new method based on flow-induced dispersion analysis to study interaction kinetics while keeping biomolecules in solution, eliminating the need for surface immobilization and thereby preserving molecular mobility and avoiding structural constraints. The method examines reactions outside equilibrium conditions by inducing a rapid concentration change in one of the binding partners (C-Jump) in a controlled microfluidic environment. Notably, it operates without buffer restrictions and requires only minimal sample quantities. We demonstrate C-Jump’s capability by accurately determining the association and dissociation rates of both protein-protein and protein-small molecule interactions. Furthermore, we validate its robustness by measuring the rates of a protein-protein interaction in human serum as well as a protein-small molecule interaction in-solution and label free. This under-lines C-Jump’s broad applicability for studying biomolecular interactions under native conditions, offering a powerful tool for advancing protein engineering and drug discovery, as well as enabling the characterization of previously inaccessible interactions.

**TABLE OF CONTENT:** This work presents C-Jump, a breakthrough method for the direct measurement of kinetic rates in solution, closely mimicking physiological conditions. Utilizing flow-induced dispersion analysis, C-Jump determines rate constants with high accuracy for protein–protein and protein– small molecule interactions. Free from buffer constraints and requiring only nanograms of protein, it can operate label-free, transforming biomolecular interaction studies across biophysics, chemistry, and medicine.

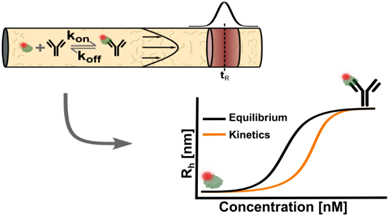

## INTRODUCTION

The ability to accurately quantify biomolecular interactions is fundamental to fields ranging from biophysics and structural biology to drug discovery and protein engineering. Kinetic measurements of protein-protein and protein-ligand interactions provide essential insights into molecular recognition, enzymatic activity, and therapeutic efficacy.^1–3^ Although several techniques exist to quantify kinetic parameters, surface-based techniques such as surface plasmon resonance (SPR) and bio-layer interferometry (BLI) have emerged as gold standards.^4,5^ However, these approaches inherently rely on immobilized binding partners, introducing potential artifacts due to surface effects, steric hindrance, and diffusion limitations.^6^ Consequently, the observed kinetics may not fully reflect interactions in solution, where biological processes naturally occur.

In-solution methods such as stopped-flow spectroscopy and kinetic isothermal titration calorimetry (ITC) have long been established for characterizing biomolecular interaction kinetics without the need for surface immobilization. Stopped-flow spectroscopy rapidly mixes two binding partners in a narrow flow channel and stops the flow to monitor changes in absorbance or fluorescence. This setup enables millisecond temporal resolution, making the method particularly suitable for fast reactions such as enzyme-catalyzed processes.^7,8^ Kinetic ITC on the other hand, determines both kinetic and thermodynamic parameters by measuring heat release during a titration, providing a label-free approach in solution.^9,10^ However, both techniques typically require sample volumes in the tens to hundreds of microliters and are not readily compatible with automated sample handling. These limitations have restricted their broader application in modern drug discovery workflows, which increasingly demand low sample consumption, automation and scalability.

Here, we introduce C-Jump, a method based on flow induced dispersion analysis (FIDA)^11,12^, that eliminates the need for surface immobilization and operates without buffer constraints, requiring only minimal protein sample. Because of its in-solution characteristics, C-Jump preserves molecular mobility, avoids structural constraints, and thus better reflects the native environment in which binding interactions typically occur. In brief, a small volume of protein sample is pulse-injected into a narrow capillary containing its binding partner. A laminar flow rapidly mixes and dilutes the sample with its binding partner inducing a concentration jump.^13^ By changing the flow velocity, the time to detection can be adjusted and, consequently, the reaction time. This allows to study the interaction before equilibrium is reached. A series of measurements at different ligand concentrations creates a binding curve outside of equilibrium, allowing precise determination of both the association rate (*k*_*on*_) and dissociation rate (*k*_*off*_).

To demonstrate C-Jump and its versatility, we apply it to measure both protein-protein and protein-small molecule interactions in solution, showing that it yields highly accurate association and dissociation rates. Furthermore, we validate its robustness with a kinetic measurement in human serum and a label-free measurement, closely mimicking true native conditions. By eliminating the constraints of immobilization-based methods, C-Jump enables kinetic analyses in solution, unlocking interactions previously inaccessible to study. Its broad applicability, ease of implementation, and compatibility with complex biological samples makes it a versatile tool for research at the interface of biophysics, chemistry, and medicine.

## RESULTS AND DISCUSSION

A non-covalent interaction between a target and a ligand (termed indicator and analyte, respectively) is governed by its association rate (*k*_*on*_) and dissociation rate (*k*_*off*_). At equilibrium, the ratio (*k*_*off*_/*k*_*on*_) defines the dissociation constant (*K*_*d*_), which characterizes the overall affinity between interaction partners. Here we present C-Jump, a method that enables precise kinetic measurements by inducing rapid concentration changes in a controlled microfluidic environment. We first outline its theoretical framework before demonstrating its applicability to three well-characterized model systems. Finally, we showcase its capability to measure in-solution kinetics of a protein-protein interaction in human serum and a label-free protein-small molecule interaction.

### C-Jump

To measure the reaction rates with FIDA under Taylor conditions^14^, two binding curves must be obtained: one at equilibrium and one outside equilibrium (Figure 1A). While equilibrium measurements are obtained from a preincubated mixture of the interaction partners (Figure1 B, top), interactions outside equilibrium are captured using the capillary mixing method (CapMix)^15^ which allows a precise control of the interaction time. In brief, a small volume of the indicator is injected between two zones of analyte. Applying a run pressure *p*_*run*_ induces a laminar flow in the capillary that rapidly mixes the components (Figure1 B, bottom). This results in a rapid change of concentration (C-Jump).^13^ In a CapMix experiment the reaction begins upon applying the run pressure *p*_*run*_. The reaction time coincides with the reference time (*t*_*R*_), which is the time it takes for the sample to reach the point of detection and is measured using the peak of the recorded fluorescence signal. By increasing *p*_*run*_, the time to detection – and thus the reaction time – can be significantly reduced (Figure1 C). Unlike traditional real-time binding assays that monitor association or dissociation continuously, C-Jump measures the apparent hydrodynamic radius *(R*_*h*_) at a defined reaction time after mixing. Each data point reflects the binding state after a specific interval, not a complete time trace of the interaction. Since 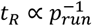, this inverse relationship enables measurements to be taken before equilibrium is reached. A detailed simulation with exact values for *t*_*R*_ and *p*_*run*_ is provided in Figure S2. To ensure measurements remain within Taylor conditions, the capillary diameter and length can be adjusted. Under these conditions, the apparent *R*_*h*_ reflects the weighted average between the diffusivity of the unbound indicator and the indicator-analyte complex.^11^ Thus, the shorter the interaction time, the smaller the apparent *R*_*h*_. This dependence of the apparent *R*_*h*_ on the interaction time provides direct kinetic insights. To quantitatively describe this relationship, we present an analytical expression that directly links the apparent *R*_*h*_ to *K*_*d*_, as well as *k*_*on*_ and *k*_*off*_ (Equation S16). Furthermore, we extend this approach to ratiometric fluorescence^16^ as an additional readout for systems where the analyte is a small molecule compared to the indicator and hence, size changes are difficult to detect (Equation S24).^17^ These expressions enable precise extraction of kinetic parameters from experimental data, allowing robust characterization of biomolecular interactions in solution under native conditions.

**Figure 1.**
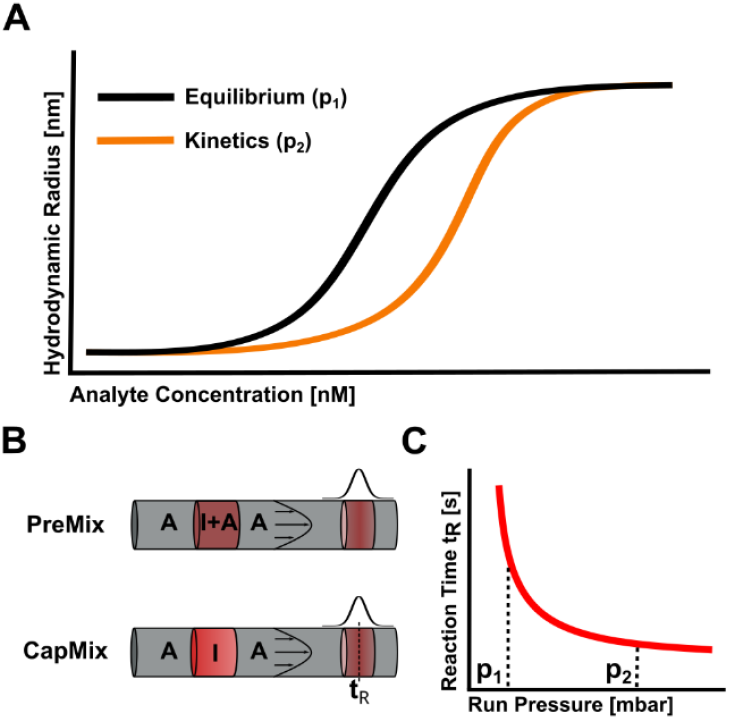
Working principle of C-Jump. (A) Sketch of binding curves under equilibrium conditions (black) and kinetic conditions (orange). Near the K_d_, the same analyte concentration results in a smaller apparent R_h_. (B) Experimental layout for PreMix and CapMix methods to study interactions at and out of equilibrium. (C) Reciprocal relationship between run pressure (p_run_) and reaction time (t_R_).

### Proof of principle

To further illustrate the principle of C-Jump, we simulated the temporal progression of complex formation for an artificial model system (Supporting Information S4) over a range of different reaction times using the analytical expression derived in Equation S16. The chosen analyte concentrations span a range centered around the *K*_*d*_, covering approximately one order of magnitude above and below to ensure enough data points in the unbound and bound regimes. As shown in Figure 2A, shorter reaction times yield smaller apparent *R*_*h*_ due to incomplete binding, with curves gradually converging toward the equilibrium profile as the reaction time increases. The inset highlights this progression for a fixed analyte concentration near the dissociation constant of the model system. To assess the robustness of the method against flow-rate variation, we further simulated datasets across a range of reaction times. We incorporated a 5 % Gaussian noise on the individual hydrodynamic radii to mimic realistic measurement uncertainty. Kinetic parameters were then extracted across a range of reaction times. The fitted on-rate constants *k*_*on*_ remained consistent across conditions, demonstrating that C-Jump yields unbiased kinetic parameters independent of reaction time and flow rate (Figure S3). However, at longer reaction times, the extraction becomes less precise, as the system approaches equilibrium and the kinetic signal becomes increasingly difficult to distinguish from the steady-state response.

**Figure 2.**
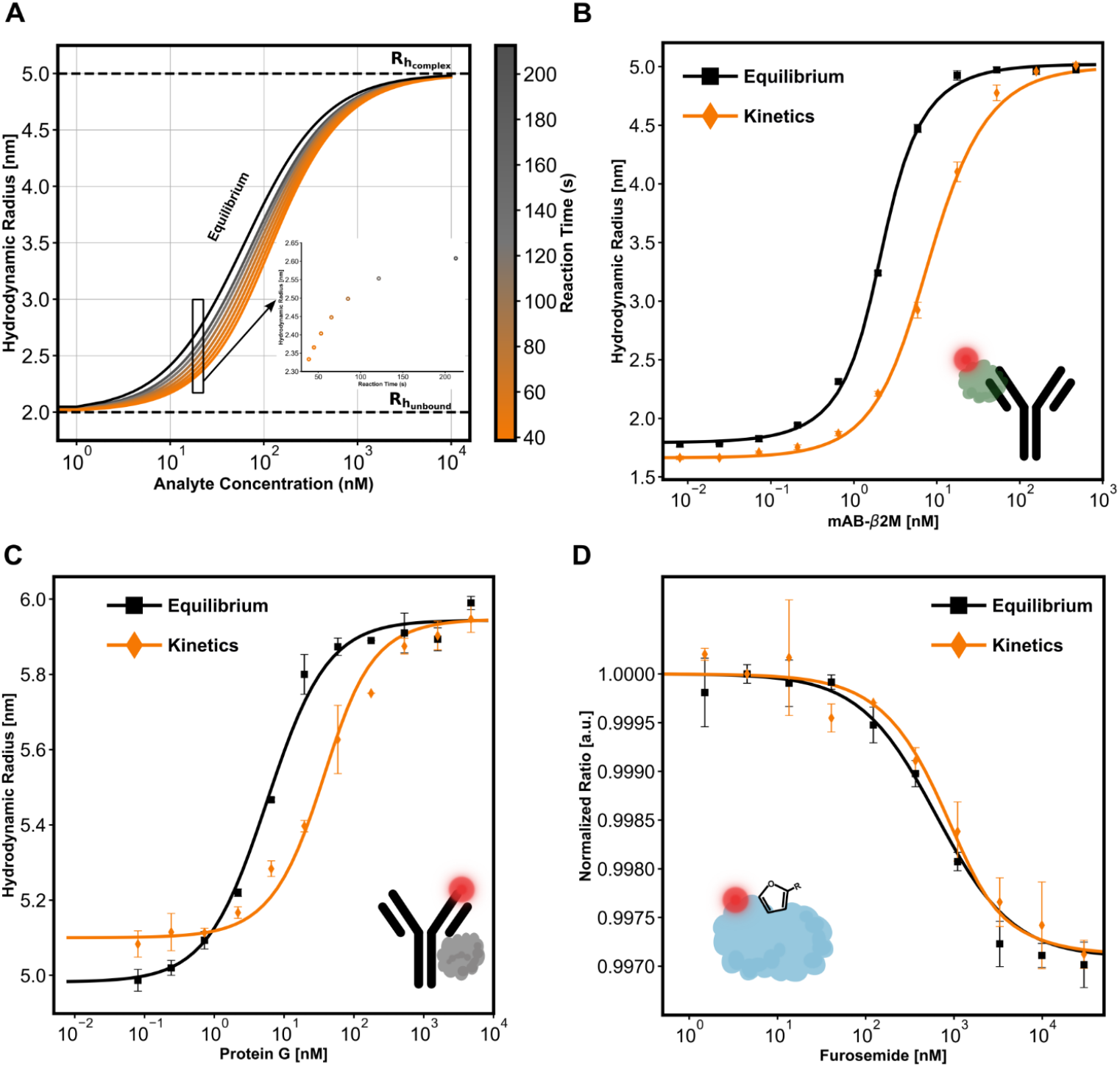
Simulation and experimental results. **(A)** Simulated apparent R_h_ as a function of analyte concentration for a range of reaction times. Curves illustrate kinetic progression from the unbound to complex state. The insert shows the apparent R_h_ versus reaction time at a fixed analyte concentration, highlighting time-resolved complex formation. **(B-D)** Measured equilibrium and kinetic binding curves for specific interactions. Each data point represents the mean of three independent measurements, with error bars indicating the standard error of the mean. **(B)** β2M* + mAb-β2M, **(C)** Rituximab* + Protein G, and **(D)** CAII* + FS. The asterix (*) marks the indicator labeled with the dye ALC647, also illustrated in the molecule sketches with a red fluorophore.

For experimental validation of C-Jump, we selected three well-characterized biomolecular interactions previously studied using surface-based methods. By comparing C-Jump data to established equilibrium and kinetic parameters, we assessed its accuracy and reliability. The resulting binding curves are shown in Figure 2B-D, with the equilibrium binding curves in black and the kinetics curves in orange. Each data point represents the mean of three independent measurements, with error bars indicating the standard error of the mean. The schematics in the corners of the graphs illustrate the respective interactions, with a red fluorophore marking the labeled indicator.

Figure 2B presents the binding curves for the protein-protein interactions of β2-Microglobulin (β2M) with its corresponding monoclonal antibody (mAB-β2M) and Figure 2C shows Rituximab with Protein G. Both interactions exhibit a measurable increase in hydrodynamic radius upon binding, with a distinct rightward shift of the kinetic curve relative to the equilibrium curve. In both cases, one of the binding partners is an antibody, with Rituximab serving as the labeled indicator and mAb-β2M as the analyte in the respective interactions. In the case of Rituximab, the apparent *R*_*h*_ of the unbound indicator corresponds to the size of an IgG antibody (∼150 kDa), whereas for β2M, the complex size also aligns with that of an IgG due to the much smaller size of β2M. Figure 2D shows the normalized binding curves for the interaction between carbonic anhydrase II (CAII) and furosemide (FS), where changes in fluorescence ratio were used to resolve binding. Despite the small relative fluorescence change (0.5 %), the binding curves remain well-defined, demonstrating the sensitivity of the detection method.

The extracted interaction parameters are summarized in Table 1, alongside values obtained from surface-based measurements reported in the literature. Minor variations, such as those observed in the CAII–FS interaction, can be attributed to differences in buffer conditions and batch-to-batch variability of the proteins. The measured value also agrees well with reported values for similar small molecule inhibitors measured with kinetic ITC.^18^ However, more substantial discrepancies, as seen in the Rituximab–Protein G interaction, are particularly evident in the dissociation rate, which is an order of magnitude faster in solution, leading to a higher *K*_*d*_. This discrepancy suggests that immobilization in surface-based assays may stabilize the complex, possibly due to avidity effects, thereby reducing its off-rate. For the interaction between β2M and mAB-β2M, the in-solution method yields a similar dissociation rate to the surface-based approach but reveals an association rate that is an order of magnitude faster. This substantial increase likely arises from the greater conformational freedom in solution, allowing the proteins to adopt favorable orientations for binding, in contrast to the restricted geometry that may be imposed by surface immobilization. Additionally, different immobilization strategies can influence affinity in surface-based assays, as demonstrated for β2M, where a four-fold change in *K*_*d*_ and two-fold variations in kinetic rates have been reported.^19^ Such effects highlight the importance of measuring interactions in solution, which reflects closer the native environment in which binding interactions typically occur.

**Table 1.** Binding affinities and kinetic rate constants measured in-solution with C-Jump and reported values for surface-based assays (asterix (*) indicates the labeled species).

| | $\beta 2M^* - mAB-\beta 2M$ | | Rituximab* – Protein G | | CAII* - FS | |
| --- | --- | --- | --- | --- | --- | --- |
|  | C-Jump | SPR<sup>19</sup> | C-Jump | SPR<sup>20</sup> | C-Jump | SPR<sup>21</sup> |
| $K_d$ [nM] | $0.32 \pm 0.05$ | 1.0 | $4.7 \pm 0.6$ | 0.9 | $555 \pm 80$ | 330 |
| $k_{on}$ [$s^{-1}M^{-1}$] | $(1.2 \pm 0.1) \times 10^7$ | $1.6 \times 10^6$ | $(2.8 \pm 0.5) \times 10^5$ | $1.9 \times 10^5$ | $(5.4 \pm 1.6) \times 10^4$ | $9.6 \times 10^4$ |
| $k_{off}$ [$s^{-1}$] | $(4.0 \pm 0.2) \times 10^{-3}$ | $1.6 \times 10^{-3}$ | $(1.2 \pm 0.2) \times 10^{-3}$ | $1.7 \times 10^{-4}$ | $(3.0 \pm 0.9) \times 10^{-2}$ | $5.0 \times 10^{-2}$ |

A key consideration of the method is the rate of complex formation, which is dependent on the concentration of interacting species. The applicability of the method can be assessed by calculating the half-life 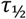 (Equation S17). For the interactions listed in Table 1, the measured half-lives range from 167 seconds for Rituximab to 7 seconds for CAII, which is the fastest half-live we measured. Faster half-lives would not be feasible to distinguish from an equilibrium measurement in the current experimental setup because the complex would be formed to a large extend before reaching the detection window. To explore this limit, we simulated binding curves for a model antibody system across a range of half-lives (5–100 seconds), showing that kinetic resolution becomes increasingly difficult below approximately 5 seconds (Figure S4). However, it can be envisioned that the experiment can be optimized to longer or shorter half-lives by adjusting the run pressure to modulate the reaction time, and the capillary dimensions to obey Taylor conditions.

### Application

We further demonstrate the capabilities of C-Jump by applying the method to study the interaction between Interleukin-2 (IL-2) and its receptor Interleukin-2Rα (IL-2Rα) under physiologically relevant conditions in 90 % human serum. This environment, which closely mimics in vivo conditions, is challenging conventional surface-based methods due to nonspecific adsorption and signal interference.^22^ To accommodate the expected fast kinetics of the interaction^23^, the experiment was performed in a shorter capillary at 15 °C. Figure 3A presents the resulting binding curves, yielding an association rate of *k_on_* = (1.9 ± 0.8) × 10^6^ M−1s−1 and a dissociation rate of *k_off_* = 0.04 ± 0.01 s−1, revealing faster kinetics compared to phosphate-buffered saline (PBS) at the same temperature (Figure S5A). Notably, the association and dissociation rates in serum at 15 °C closely matched those measured in PBS at 25 °C (Table S2), as well as previously reported values from a surface-based assay.^23^ A potential explanation for this acceleration is a structural rearrangement of IL-2 in serum, as indicated by the significantly larger apparent *R*_*h*_ of both the indicator and the complex compared to their sizes in PBS (Figure S6). Similar serum-induced structural changes have been reported for other proteins^24^ and may expose the binding pocket, facilitating faster binding and unbinding events. Interestingly, direct measurements in serum at 25 °C were not feasible, as the rapid kinetics exceeded the experimental detection limit. However, given that the kinetics in serum at 15 °C align with those in PBS at 25 °C, it is likely that the interaction in serum at 25 °C follows a similar trend. These findings emphasize the importance of performing kinetic measurements in truly native biological environments, as serum-specific structural effects can significantly influence interaction dynamics, which may be overlooked in simplified buffer conditions.

**Figure 3.**
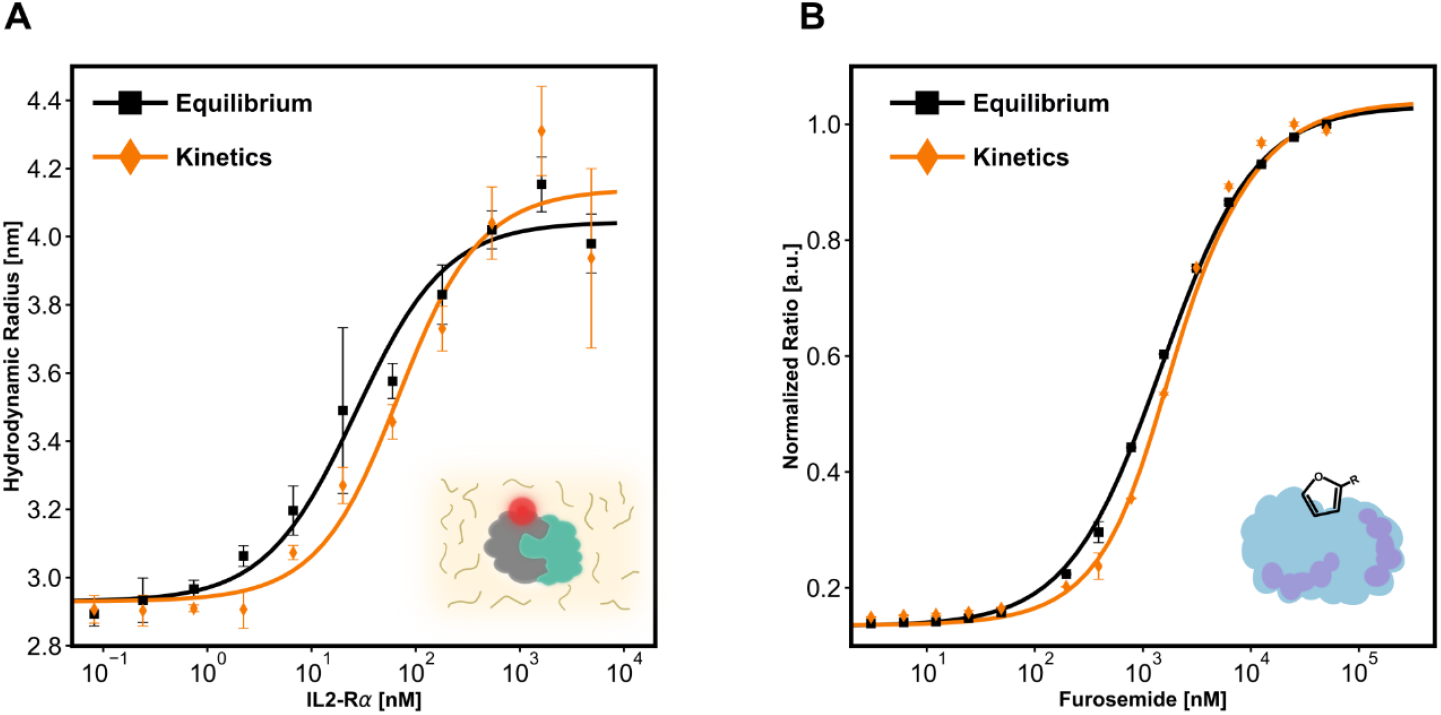
Measured equilibrium and kinetic binding curves in human serum and label-free. Each data point represents the mean of three independent measurements, with error bars indicating the standard error of the mean. **(A)** Kinetics of IL-2 and its receptor IL-2Rα in 90 % human serum. The K_d_ is 19 nM with the rate constants k_on_ = (1.9 ± 0.8) × 10^6^ M^™1^s^™1^ and k_off_ = 0.04 ± 0.01 s−1. The apparent R_h_ of the unbound indicator and the complex are significantly bigger than measured in pure buffer (Figure S6). **(B)** Kinetics of CAII binding to FS, measured in solution and label-free using ratiometric fluorescence of tryptophan residues. The K_d_ is 402 nM with the rate constants k_on_ = (1.8 ± 0.2) × 10^5^ M−1s−1 and k_off_ = 0.07 ± 0.01 s−1.

To further demonstrate the potential of C-Jump, we performed a label-free, in-solution measurement of the interaction between CAII and FS. We leveraged the ratiometric fluorescence of tryptophan residues (Supporting Information S7), which are highly sensitive to their local environment.^25,26^ As shown in Figure 3B, the resulting binding curves yield an association rate of *k_on_* = (1.8 ± 0.2) × 10^5^ M−1s−1 and a dissociation rate of *k_off_* = 0.07 ± 0.01 s−1, representing a similar result compared to the values reported in Table 1. The close agreement between labeled and label-free results supports that labeling had minimal impact under the conditions used. Notably, the fluorescence ratio exhibited a remarkable 900 % relative change, corresponding to an 1800-fold increase in sensitivity compared to the labeled experiment. This highlights the exceptional sensitivity of the detection method and demonstrates the potential of C-Jump for studying proteins close to their native condition.

## CONCLUSION

Accurate quantification of biomolecular interactions is essential across a broad spectrum of disciplines, including biophysics, structural biology, drug discovery, and protein engineering. Although numerous methods exist to determine binding affinity, the measurement of kinetic rates is predominantly limited to surface-based techniques, which rely on immobilized binding partners and may not accurately mimicking physiological conditions. In this work, we introduce C-Jump, a paradigm shift that directly measures kinetic rates in solution, more closely reflecting the native environment where binding typically occurs. Leveraging flow-induced dispersion analysis, C-Jump is free from buffer constraints, can operate label-free and requires only nanograms of protein. These unique features make C-Jump a versatile and efficient approach, positioning it as a transformative tool for advancing research in drug discovery, biophysics, chemistry, and medicine.

## MATERIALS & METHODS

### Equipment

The experiments were conducted using a Fida Neo 640 system (Catalog No. N002-640, Fida Biosystems ApS) and a Fida Neo UV detector (Catalog No. Det-UVN, Fida Biosystems ApS). Experiments utilize two types of fused silica capillaries (Ø 75 µm, L = 100 cm, Catalog No. 100-001 and Ø 50 µm, L = 70 cm, Catalog No. 100-057, Fida Biosystems ApS). Capillaries were pre-treated by flushing with a high-sensitivity coating reagent (Catalog No. 310-010, Fida Biosystems ApS) prior to each experiment.

### Materials and Chemicals

Human β2-Microglobulin (B2M-H5225, Lot 475-2391F1-1LQ), human Interleukin-2 (IL2-H5215) and human Interleukin-2Ralpha (ILA-H52H9) were purchased from Acro biosystems. Anti-β2-Microglobulin monoclonal antibody was sourced from Cytiva (28980886). Bovine carbonic anhydrase II (C2522-1G, Lot No. SLCQ8559) and furosemide (F4381, Lot No. MKCM6678) purchased from Sigma-Aldrich. Rituximab (MA5-47773) and Protein G (101201) were purchased from Thermo Fisher. Phosphate-buffered saline (PBS) 1x was prepared from a 10x PBS stock solution (Lot No. Q01I511, Thermo Scientific) using deionized water (Direct-Q3, Sigma-Aldrich). Additional reagents included human serum type AB (male, H4522-20ML, Sigma-Aldrich) and sodium hydroxide 1 M (NaOH) from sodium hydroxide pellets (Lot No. SLCC5273, Sigma-Aldrich).

### Assay buffer

The assay buffer for all binding studies is PBS (1x, pH 7.4) supplemented with 0.03 % Pluronic F-127 (Millipore, Lot No. 3951203) to prevent adsorption in the 96-well plates.

### Protein Labeling

The storage buffer for β2-Microglobulin (β2M) was exchanged with PBS (1x, pH 7.4) using a spin column (Catalog No. 89882, Thermo Scientific), which was pre-rinsed three times with PBS at 1.5 × 1000 rpm for 60 seconds per rinse. After buffer exchange, the protein concentration was adjusted to 1 mg/mL. Carbonic anhydrase II (CAII) was dissolved in PBS to final concentrations of 1 mg/mL. Lyophilized Interleukin-2 (IL-2) was dissolved in deionized water to a concentration of 1 mg/ml. Proteins were individually labeled using the Fida 1 Protein Labeling Kit ALC 640 (Cat. No. 430-003, Fida Biosystems ApS). Labeling was achieved via a Cy5 related, conjugated NHS ester dye, which reacts predominantly with primary amino groups, to form stable amide bonds. Proteins were diluted to approximately 1 mg/mL in PBS and mixed with 1 M sodium bicarbonate buffer (pH ∼8.3). The dye was then added at a fourfold molar excess. The reaction was incubated in the dark for 30 minutes at room temperature. Unreacted dye was removed using a spin column purification step provided in the kit. Degrees of labelling were 1.8 for β2M, 1.4 for CAII, 0.9 for Rituximab and 0.1 for IL-2 (NanoDrop, Thermo Fisher). Free dye content for all proteins was less than 5 % measured with a Fida Neo system 640. For the measurement in human serum, any remaining free dye was removed before the experiment with a spin column provided in the labelling kit. Labeled proteins, in the following referred to as indicator, were stored at - 20 °C.

### Ligand Preparation

**\**Anti-β2-Microglobulin monoclonal antibody (mAB-β2M) was received at a stock concentration of 1 mg/mL and stored at 4 °C. Stock solution of the CAII inhibitor furosemide (FS) was prepared in DMSO at 100 mM and stored at 4 °C. Protein G and Interleukin-2Ralpha (IL-2Rα) were prepared from lyophilized powder by adding deionized water to a concentration of 1 mg/mL and stored at -20 °C.

### Premix Experiment

Stock solutions of the ligand were diluted in PBS or serum to the following concentrations: 1 µM for mAB-β2M, 10 µM for Protein G, 50 µM for FS, and 10 µM for IL-2Rα. A 12-point dilution series (3-fold dilution) was prepared for each analyte in the respective buffers. A portion of the analyte was transferred and premixed with the corresponding indicator to achieve a final volume of approximately 20 µL. The indicator concentrations for the indicators were 10 nM for β2M, 10 nM for Rituximab, 100 nM for CAII, and 30 nM for IL-2. The indicator concentration for the unlabeled CAII was 600 nM.

Unless stated otherwise in the text, all experiments were performed at 25 °C. Each data point required 40 nL of indicator and 12 µL of analyte and took approx. 5 min. The experimental sequence included: (1) flushing with NaOH at 3500 mbar for 20 s; (2) flushing with deionized water at 3500 mbar for 20 s; (3) flushing with high-sensitivity coating reagent (amphiphilic block polymer, passivating the surface and making it hydrophilic) at 3500 mbar for 20 s. (4) flushing with deionized water at 3500 mbar for 20 s. (5) filling the capillary with analyte solution at 3500 mbar for 20 s; (6) injecting premixed-indicator at 50 mbar for 10 s; (7) mobilizing and measuring with analyte solution at 400 mbar for 180 s. The signal in both spectral channels was recorded simultaneously for the measurement time during step 7.

### Kinetics Experiment

The kinetics experiments were performed immediately following the equilibrium experiment, using the same dilution series of the analyte. The indicator concentration was identical to that used in the premix and was prepared separately in the experimental buffer. For the serum experiment, the indicator was kept in PBS to prevent degradation during the experiment.

The procedure for step 1) to 5) is equal to the equilibrium experiment. (6) injecting unbound indicator at 50 mbar for 10 s; (7) mobilizing and measuring with analyte solution at pressures and times as shown in Table 2, including also the used capillary in diameter × length. The signal in both spectral channels was recorded simultaneously for the measurement time during step 7.

**Table 2.** Conditions for kinetics experiments.

| Interaction | Run Pressure [mbar] | Time [s] | Capillary [$\mu\text{m} \times \text{m}$] |
| --- | --- | --- | --- |
| $\beta 2\text{M}$ & mAB-$\beta 2\text{M}$ | 1000 | 50 | $\varnothing 75 \times \text{L1}$ |
| Rituximab & Prot. G | 400 | 60 | $\varnothing 75 \times \text{L1}$ |
| CAII & FS | 1600 | 45 | $\varnothing 75 \times \text{L1}$ |
| CAII & FS (UV) | 800 | 45 | $\varnothing 75 \times \text{L0.7}$ |
| IL-2 & IL-2Rα | 3000 | 60 | $\varnothing 50 \times \text{L0.7}$ |

### Data Processing

Data analysis was performed using FIDA Analysis Software (Version 3.1.1.0, Fida Biosystems ApS). Multi-species Gaussian fitting was applied to distinguish between the signal from the labeled protein and the free fluorophore. The extracted peak width of the labeled protein was used to determine its apparent hydrodynamic radius.

For ratiometric fluorescence measurements, the fitting procedure was applied to both spectral channels, with the area under the curve representing the total fluorescence. The ratio was then calculated by dividing the total fluorescence of the longer-wavelength spectral channel by that of the shorter-wavelength spectral channel. The mean and standard error of the mean were plotted against the analyte concentration. Binding isotherms were then fitted to obtain dissociation constants, while kinetic experiments provided reaction rate constants. Detailed descriptions of the applied models can be found in the supporting information S1 and S5.

## ASSOCIATED CONTENT

### Supporting Information

Additional experimental details, data analysis, materials and methods, including schematics of experimental setup (PDF).

## Acknowledgement

Financial support from the Innovation Fund Denmark (grant no. 2052-00002B) is gratefully acknowledged.

