## supplementary file for "In-Solution Characterization of Biomolecular Interaction Kinetics under Native Conditions"

*Henrik Jensen*[1,*]

[1]FIDA Biosystems ApS, 2860 Søborg, Denmark

[2]Technical University of Denmark, Department of Health Technology, 2800 Kongens Lyngby, Denmark

**Table of Contents**


S1 – Theoretical derivation of the C-jump model based on the diffusivity S3

S2 – Dilution factor γ S9

S3 – Simulation of Reaction Times S11

S4 – Simulation of C-Jump S12

S5 – C-Jump based on ratiometric fluorescence S15

S6 – Interleukin-2 & Interleukin-2Rα S17

S7 – Setup for ratiometric fluorescence measurement of tryptophan residues S19

References S20

### S1 – Theoretical derivation of the C-jump model based on the diffusivity

*Table S1. Parameters and corresponding units used in the analytical derivation.*

| Symbol | Description | Unit |
|---|---|---|
| $K_d$ | Equilibrium dissociation constant | M |
| $k_{off}$ | Dissociation rate constant | $s^{-1}$ |
| $k_{on}$ | Association rate constant | $M^{-1}s^{-1}$ |
| $A_T$ | Total concentration of analyte | M |
| $I_T$ | Total concentration of indicator | M |
| $R_c$ | Capillary radius | m |
| $u_0$ | Maximum velocity in capillary | $m \cdot s^{-1}$ |
| $k(t)$ | Dispersion coefficient | $m^2\ s^{-1}$ |
| $D(t)$ | Diffusivity | $m^2\ s^{-1}$ |
| $I_d$ | Local indicator concentration | M |
| $\boldsymbol{\sigma}$ | Measured peak variance | s |
| $R_h$ | Apparent hydrodynamic radius | m |
| $T$ | Temperature | K |
| $k_B$ | Boltzmann constant | $J \cdot K^{-1}$ |
| $\boldsymbol{\eta}$ | Viscosity | Pa·s |

The subscripts I and IA refer to the unbound indicator and complex, respectively.

Derivation

We consider the chemical reaction between a target (further called indicator *I*) and its ligand (further called analyte *A*). The interaction between I and A can be described as

$$IA \underset{k_{on}}{\overset{k_{off}}{\rightleftharpoons}} I + A$$

*(S1)*

Where *IA* is an in-solution complex composed of *I* and *A*. The dissociation constant $K_d$ can be described by the on-rate ($k_{on}$) and off-rate ($k_{off}$) which describe the rate of association in $\mathrm{M}^{-1}\mathrm{s}^{-1}$ and the rate of dissociation in $\mathrm{s}^{-1}$, respectively.

$$K_d = \frac{k_{off}}{k_{on}} = \frac{[A][I]}{[AI]}$$

(S2)

Square brackets denote the concentrations, where we further define the total concentration of analyte $A_T = [A] + [IA]$ and indicator $I_T = [I] + [IA]$.

We consider experimental conditions where Taylor conditions are fulfilled.[1] Further, the injected *I*-zone is only a very small part of the capillary (< 1 %). Under such conditions *A* is mixed into the *I*-zone rapidly and it can be assumed that *A* is uniformly distributed in the capillary at all times $t > 0$. In this case, the apparent diffusion coefficient describing the dispersion is a time dependent weighted average between the diffusivity of bound and unbound *I*.

Given this experimental framework, the mass transport problem can be simplified and described as[1]

$$\frac{\partial c(x,t)}{\partial t} = k(t)\frac{\partial^2 c(x,t)}{\partial x^2}$$

(S3)

Where $k(t)$ is the dispersion coefficient which is linked to the time dependent diffusivity $D(t)$ and the capillary radius $R_c$ and the maximum velocity in the center of the capillary $u_0$ according to:

$$k(t) = \frac{R_c^2 u_o^2}{192} \cdot (1/D(t))$$

(S4)

In the limiting case of a constant diffusivity (i.e. at equilibrium), $k(t) = const$ . The factor 192 in the nominator arises from the analytical solution of Taylor dispersion in a circular tube.[1]

Here, we model the diffusivity as weighted average of bound and unbound indicator which is time dependent (kinetics of interaction) but decoupled from spatial diffusion. Therefore, we introduce a convoluted diffusivity $\tau_D$ , which can be seen as an average dispersity in an experiment:

$$\tau_{D1} = \int_0^{t_R} k(t)dt = \widehat{k1} \cdot \int_0^{t_R} \frac{1}{D(t)} dt$$

(S5)

Where

$$\widehat{k1} = \frac{R_c^2 u_o^2}{192}$$

Hence, by change of variables Equation (S3) can be expressed as

$$\frac{\partial c(x, \tau_{D1})}{\partial \tau_{D1}} = \frac{\partial^2 c(x, \tau_{D1})}{\partial x^2}$$

(S6)

Due to the same form of the equation, a general Gaussian solution can be found in the same way considering Taylor condition and boundary conditions for pulse injection given by Taylor at a point $x_1$:[1]

$$c(x_1, \tau_{D1}) = k_1 \cdot e^{-\frac{(x_1)^2}{4 \cdot \tau_{D1}}}$$

(S7)

The special position $x_1$ can be converted into a time trace using:

$$(x_1) \quad = 1/2(u_0) \quad (t - t_r)$$

$$(x_1)^2 = 1/4(u_0)^2(t - t_r)^2$$

Where ½$u_0$ is the average speed of the plug in the capillary. Thus, the concentration can be written as:

$$c(t) = k_1 \cdot e^{-\frac{(t-t_r)^2}{4\tau_D}}$$

(S8)

Where

$$\tau_D = \hat{k} \cdot \int_0^{t_R} \frac{1}{D(t)} dt$$

(S9)

with

$$\hat{k} = \frac{R_c^2}{48}$$

The factor 48 in the numerator results from algebraic rearrangement of the Taylor dispersion correction term, originally expressed above with a denominator of 192. Then, $\tau_D$ is related to the measured peak variance $\sigma^2$ at the reference time $t_R$ according to(S10)**Error! Reference source not found.**

$$\tau_D = ½\sigma^2$$

*(S10)*

Where $\sigma^2$ can be obtained from fitting the raw fluorescence data $S(t)$ with Equation (S11). The fit yields the reference time $t_R$, the constant background offset $C_1$, the response factor $C_2$ and the width of peak $\sigma$.

$$S(t) = C_1 + C_2 \cdot exp\left[-\frac{(t - t_R)^2}{2\sigma^2}\right]$$

*(S11)*

Having now established a value for $\tau_D$, we look closer at the apparent diffusivity $D(t)$, which changes according to the fraction bound of *I* ($x = [IA] \cdot ([I] + [IA])^{-1}$). At the start, no *I* is bound to *A* but it increases during the time of the experiment. Therefore, the diffusivity can be expressed as

$$D(t) = D_{IA}x(t) + D_I(1 - x(t))$$

*(S12)*

The concentration of formed complex *[IA]* at time *t* is given by the solved rate equation (S1) with boundary condition $[IA](t = 0) = 0$. We introduce $I_d = \gamma I_T$ which is the dispersion-induced local concentration of *I* in the dispersion zone at $t_R$, i.e. $I_d = I_t(t = t_R)$. It is related to the total concentration through the dilution factor $\gamma$ (Equation (S21)).

$$[IA](t) = \frac{I_d \cdot A_T}{A_T + I_d + K_D} \cdot \left(1 - e^{-(k_{off} + k_{on} \cdot (A_T + I_d)) \cdot t}\right)$$

*(S13)*

We combine Equation (S12) and Equation (S13) through the above given expression for *x*:

$$D(t) = D_I + (D_{IA} - D_I) \cdot \frac{A_T}{A_T + I_d + K_d} + (D_I - D_{IA}) \cdot \frac{A_T}{A_T + I_d + K_d} \cdot e^{-(k_{off} + k_{on} \cdot (A_T + I_d)) \cdot t}$$

*(S14)*

In the limiting case of $t \to \infty$, we approach the equilibrium state of a simple 1:1 interaction and the Equation (S14), simplifies to the first two terms, corresponding to the FIDA binding isotherm presented by Jensen et al.[2]

For easier bookkeeping we rewrite Equation (S14) to its final form with the new parameters *p, q* and *a*:

$$D(t) = p + qe^{at}$$

$$p = D_I + (D_{IA} - D_I) \cdot \frac{A_T}{A_T + I_d + K_d}$$

$$q = (D_I - D_{IA}) \cdot \frac{A_T}{A_T + I_d + K_d}$$

$$a = -\left(k_{off} + k_{on} \cdot (A_T + I_d)\right)$$

After obtaining this expression for *D(t)*, we can plug it back into Equation (S9) and evaluate the integral at $t_R$ to obtain $\tau_D$:

$$\tau_D = \int_0^{t_R} k(t)dt = \hat{k} \cdot \int_0^{t_R} \frac{1}{D(t)} dt = \hat{k} \cdot \int_0^{t_R} \frac{1}{p + q \cdot e^{a \cdot t}} dt$$
$$= \frac{\hat{k}}{p} \cdot \left\{ t_R + \frac{1}{a} \cdot ln \left[ \frac{p + q}{p + q \cdot e^{a \cdot t_R}} \right] \right\}$$

*(S15)*

In an actual kinetics experiment, the apparent diffusivity ($D_{app}$) obtained from a single species fit to the raw data can be obtained as:

$$D_{app} = \frac{R_c^2 \cdot t_R}{24 \cdot \sigma^2}$$

Lastly, $\tau_D$, can be linked to the hydrodynamic radius $R_h$ through the Stokes-Einstein relation according to

$$R_h = \frac{k_B \cdot T}{6 \cdot \pi \cdot \eta \cdot D_{app}} = \frac{k_B \cdot T \cdot 24 \cdot \sigma^2}{6 \cdot \pi \cdot \eta \cdot R_c^2 \cdot t_R} = \frac{k_B \cdot T \cdot 24 \cdot 2 \cdot \tau_D}{6 \cdot \pi \cdot \eta \cdot R_c^2 \cdot t_R}$$

$$R_h = \frac{8 \cdot k_B \cdot T \cdot \tau_D}{\pi \cdot \eta \cdot R_c^2 \cdot t_R}$$

*(S16)*

This links $\tau_D$ to $R_{h,I}$, $R_{h,IA}$, $K_d$ and the kinetics ($k_{on}$ and $k_{off}$). Hence, measuring a series of $\tau_D$ in a titration of varying $A_T$, for different time points $t_R$ (achieved by varying the run-pressure) allows to fit Equation (S16), and thus, to extract the kinetics of the interaction. However, an easier way is to first do an experiment at equilibrium, to obtain $R_{h,I}$, $R_{h,IA}$, $K_d$ and afterwards an additional experiment outside the equilibrium to measure $k_{on}$ and $k_{off}$.

A measure for the applicability of the method is the half-life $\tau_{½}$ of the interaction, which is concentration dependent. Evaluating the half-live at a concentration equal to Kd/2 simplifies the equation as shown in Equation (S17). We see applicability of the method for half-lives down to $\tau_{½} \approx 5\ s$.

$$\tau_{½} = \frac{\ln(2)}{k_{obs}} = \frac{\ln(2)}{k_{on} \cdot [A]_{0.5K_d} + k_{off}} = \frac{\ln(2)}{k_{on} \cdot 0.5 \cdot K_d + k_{off}} = \frac{2\ \ln(2)}{3\ k_{off}}$$

*(S17)*

### S2 – Dilution factor $\gamma$

Since the indicator zone is dispersed and thereby diluted, the local indicator concentration at time point $t_R$ can not be approximated with $I_T$. The dilution can be modelled, knowing the injection time of the indicator $t_i$ and the flow velocities $u_i$ and $u_r$ during the injection and during the actual experiment, respectively.

The initial spatial width of the indicator zone is $\delta_i = u_i t_i$. During the experiment, the temporal width of the zone can be extracted from the gaussian distribution describing the Taylor dispersion, which can be expressed as spatial width according to Equation (S18). Here, *D* is the average diffusion in the experiment, which can be obtained as described above.

$$\delta_r = u_r \cdot 2 \cdot \sigma(t) = 2u_r \sqrt{\frac{R_c^2}{24D}} \cdot \sqrt{t} \tag{S18}$$

The spatial widths $\delta_r$ and $\delta_s$ are linked through an average dilution factor $\gamma(t)$, given in Equation (S19). Using the Hagen-Poiseuille equation, we express the dilution factor in terms of injection and run pressure rather than flow velocities.

$$\gamma(t) = \frac{\delta_i}{\delta_i + \delta_r} = \frac{u_i \cdot t_i}{u_i \cdot t_i + u_r \cdot 2 \cdot \sigma(t)} = \frac{P_i \cdot t_i}{P_i \cdot t_i + P_r \cdot 2 \cdot \sigma(t)} \tag{S19}$$

We further combine Equation (S18) and Equation (S19) to obtain the expression

$$\gamma(t) = \frac{1}{1 + b \cdot \sqrt{t}}$$

$$b = 2 \cdot \sqrt{\frac{R_c^2}{24 \cdot D}} \cdot \left(\frac{P_r}{P_i \cdot t_i}\right)$$

All parameters in *b* are either known from the experiment or can be directly obtained from the experimental data. Dilution is a gradually process and we therefore assume an average dilution factor from the start of the experiment until the time of detection $t_R$ according to

$$\gamma = \frac{1}{t_R - 0}\int_0^{t_R} \gamma(t)dt = \frac{1}{t_R}\cdot\int_0^{t_R} \frac{1}{1 + b\cdot\sqrt{t}}dt = \frac{2}{b\cdot\sqrt{t_R}} - \frac{2}{b^2\cdot t_R}\cdot\ln\left(b\cdot\sqrt{t_R} + 1\right)$$

*(S20)*

Hence, the average total indicator concentration during the experiment is given by.

$$I_d = \gamma I_T$$

*(S21)*

Figure S1 shows the dilution factor for different run pressures in two different capillaries ($d = 75\ \mu m\ with\ l = 1\ m\ and\ d = 50\ \mu m\ with\ l = 0.7\ m$) length. As an example, for $P_{run} = $ 3000 mbar, the dilution factor in the thinner capillary (d= 50 μm) is 0.07 and therefore an initially injected indicator concentration of 10 nM is diluted to 0.7 nM at the point of detection.

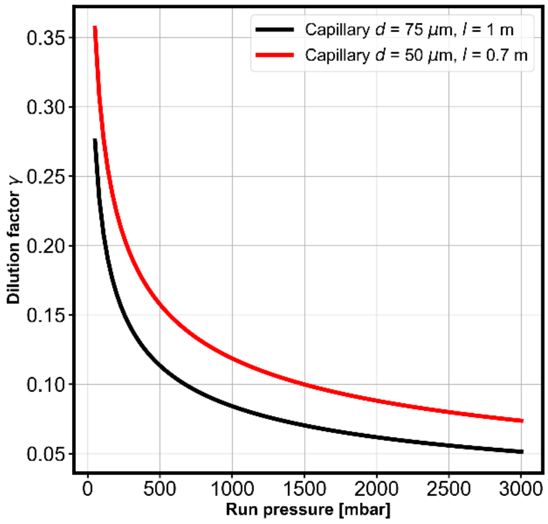


*Figure S1. Dilution factor γ versus run pressure in two different types of capillaries. For the simulation the injection pressure is $P_i = 50\ mbar$ for an injection time of $t_i = 10\ s$. The diffusivity is $D = 9.81 \times 10^{-11}\ m^2 s^{-1}$, corresponding to a hydrodynamic radius of $R_h = 2.5\ nm$.*

### S3 – Simulation of Reaction Times

The reaction time $t_R$ is simulated as function of the run pressure $p_{run}$ in a capillary of 75 µm inner diameter and 1 m length.

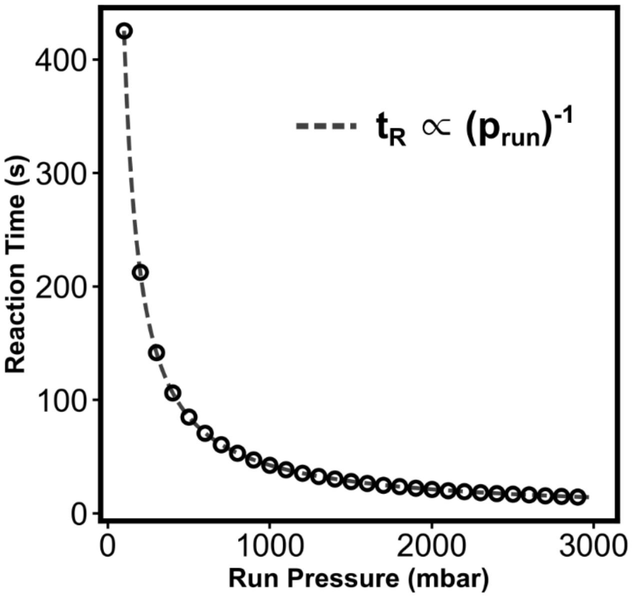


*Figure S2. Simulation of reaction time tR as function of run pressure in a capillary of 75 µm inner diameter and 1 m length. The relationship follows directly from the Hagen–Poiseuille equation and demonstrates a reciprocal dependence on pressure ($t_R \propto 1/p_{run}$). This enables precise experimental control over reaction time by simply adjusting run pressure in the C-Jump method*

### S4 – Simulation of C-Jump

To simulate the apparent $R_h$ shown in Figure 2A (main text), we used the analytical model from Equation (S16), assuming a 1:1 interaction with the following fixed parameters:

$$R_{h,I} = 2\ nm \qquad T = 25\ °C$$
$$R_{h,IA} = 5\ nm \qquad \eta = 0.00089\ Pa \cdot s$$
$$c_I = 10\ \text{nM} \qquad r_c = 37.5\ \mu\text{m}$$
$$c_A = 0\ \mu\text{M to } 10\ \mu\text{M} \qquad l_c = 1.0\ \text{m}$$
$$K_d = 25\ nM$$
$$k_{on} = 1.0 \cdot 10^6 M^{-1} s^{-1}$$
$$k_{off} = 2.5 \cdot 10^{-2} s^{-1}$$

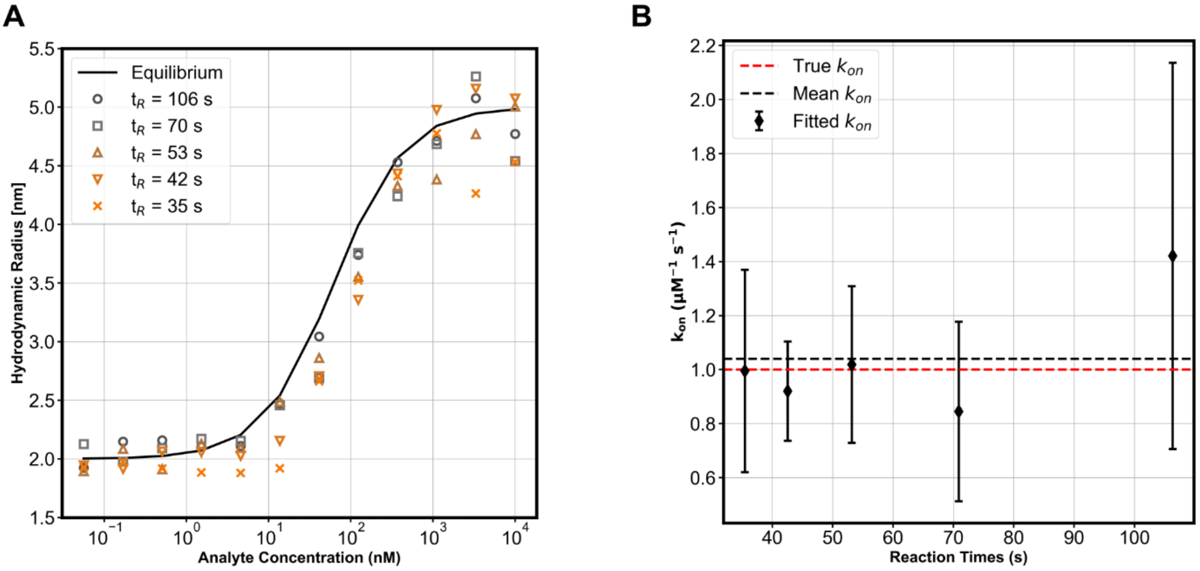


*Figure S3. Simulation of a model system to assess the robustness of the C-Jump method across different reaction times $t_R$. **(A)** The apparent hydrodynamic radii were simulated over a range of analyte concentrations (3-fold dilution, 12-points) for different reaction times (35–106 s). Gaussian noise (5 %) was added to each measurement point to mimic realistic measurement uncertainty. The solid black line represents the binding curve at equilibrium (i.e. $t_R \to \infty$). **(B)** On-rate constants $k_{on}$ were extracted from fits to the noisy simulated curves in A. The error bars resemble the standard deviation on the fit. The dashed red line marks the true $k_{on}$ value used for simulation $(1.0 \cdot 10^6 M^{-1} s^{-1})$, and the dashed black line shows the mean of all extracted values. No systematic bias is observed across reaction times. Slightly larger fitting errors at longer reaction times are expected due to reduced kinetic resolution as the system approaches equilibrium. Simulation parameters as given above.*

To further assess the lower limit of resolvable half-lives, we simulated a range of kinetic scenarios using the same model system as described above. The simulations span half-lives from 5 to 100 seconds, which correspond to $k_{off}$ values according to Equation (S17). Given a fixed $K_d$ of 25 nM, the associated $k_{on}$ values were determined using Equation (S17). The reaction time was set to the shortest duration to obey Taylor condition for a complex with a hydrodynamic radius of $R_{h_{AI}} = 5$ nm.

Two sets of capillary dimensions were simulated: one with a length of L = 1 meter and an inner diameter (ID) of ID = 75 µm, and one with a length of L = 0.7 m and ID = 75 µm. The reaction times are 35 seconds and 18 seconds, respectively.

The simulation results for these configurations are shown in Figures S4A and S4C, respectively. Figures S4B and S4D display the corresponding $k_{on}$ values extracted by fitting Equation (S16) with black diamonds representing the fitted values and error bars indicating the standard deviation of the fit. Red circles denote the ground truth values used in the simulation. As expected, shorter half-lives (i.e., faster kinetics) lead to curves that more closely resemble the equilibrium binding profile, making the kinetic parameters more difficult to resolve with precision.

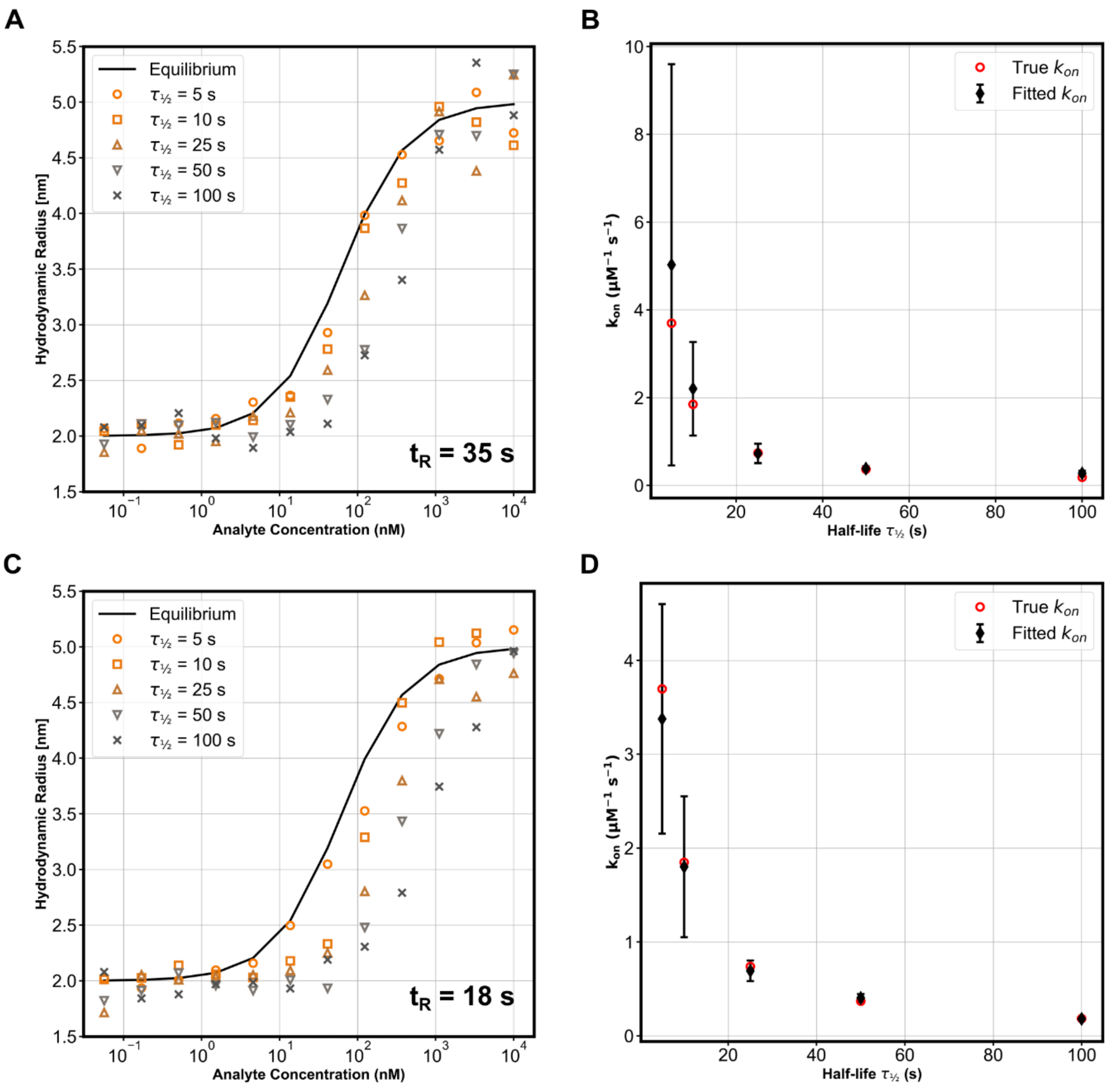


*Figure S4. Simulation of kinetic resolution limits for fast interactions with half-lives ranging from 5 to 100 s.*
***(A, C)*** *Simulated apparent $R_h$ as a function of analyte concentration for capillary lengths of 1 m (A) and 0.7 m (C), with inner diameters of 75 µm and 50 µm, respectively. Reaction times were 35 s and 18 s, respectively, corresponding to the shortest durations satisfying Taylor dispersion for a complex of size $R_{h_{AI}}$ = 5 nm.*
***(B, D)*** *Extracted $k_{on}$ values (black diamonds) and their standard deviations from fitting Equation (S16) to the simulated curves. Red circles denote the true input $k_{on}$ values derived from fixed $K_d$ = 25 nM and variable $\tau_{1/2}$. Faster interactions result in reduced resolution due to limited deviation from the equilibrium profile.*

### S5 – C-Jump based on ratiometric fluorescence

Following a similar approach, we find the analytical expression for C-Jump based on a ratiometric fluorescence measurement. The fluorescence intensity is detected in two separate spectral bands within the emission spectrum of the fluorophore, which makes it sensitive to spectral related intensity changes, often induced by small molecule binding or conformational changes.

The fluorescence intensity is quantified as the area under the curve of the detected signal *S(t)*. This area can be obtained from the same fit described in Equation (S11), where it is given by $F = C_2\sigma\sqrt{2\pi}$.

For the C-Jump analysis, the starting point is Equation (S22).[3] As before, *x* represents the fraction of bound indicator. The factors $\lambda_{IA}$ and $\lambda_I$ are the fluorescence ratios of the complex and unbound indicator, respectively. The correction factor $s = F_{u_1}F_{b_1}^{-1}$ links the fluorescence intensity of bound and unbound indicator in the first spectral band.

$$F_{2/1} = \frac{\lambda_{IA}x + s\lambda_I(1-x)}{x + s(1-x)}$$

(S22)

We use the same expression for *[AI]* as given in Equation (S13) and thus

$$x = b(1 - e^{-at})$$

$$a = k_{on}(K_d + I_d + A_t)$$

$$b = \frac{A_t}{A_t + I_d + K_d}$$

(S23)

Unlike diffusivity, fluorescence intensity does not require evaluation over the entire experiment duration (i.e. no integration from zero to $t_R$). Therefore, we can rewrite Equation (S22)(S22) and plug in Equation (S23) to obtain the final expression

$$F_{2/1} = \frac{\frac{A_t}{A_t + I_d + K_d}\left(1 - e^{-\left(k_{on}(K_d + I_d + A_t)\right)}\right)(\lambda_{IA} - s\lambda_I) + s\lambda_{IA}}{\frac{A_t}{A_t + I_d + K_d}\left(1 - e^{-\left(k_{on}(K_d + I_d + A_t)\right)}\right)(1 - s) + s}$$

(S24)

This links the fluorescence ratio to the kinetics $k_{on}$ and $k_{off}$. The easiest way to apply this equation is to do first an experiment at equilibrium condition to obtain $\lambda_{IA}$, $\lambda_I$, $s$ and $K_d$. Then, in a following experiment outside equilibrium conditions the only remaining free parameter to fit is $k_{on}$.

### S6 – Interleukin-2 & Interleukin-2Rα

The interaction between interleukin-2 (IL-2) and its receptor interleukin-2 receptor alpha (IL-2Rα) was studied in phosphate-buffered saline (PBS). A 12-point binding curve was obtained using a three-fold serial dilution of IL-2Rα, starting at a concentration of 5 μM. The experiment was conducted in a fused silica capillary (Ø 50 μm, L = 0.7 m), which was dynamically coated with a polymer before each run to prevent protein adhesion to the capillary walls. For the equilibrium binding curve, samples were pre-mixed and incubated prior to measurement. The kinetic binding curve was acquired using a CapMix experiment[4] at a run pressure of 3000 mbar to capture rapid interaction dynamics. Figure S5 presents the resulting binding curves along with the extracted dissociation constant ($K_d$) and kinetic rate constants ($k_{on}$ and $k_{off}$). The results and previously reported results from literature are tabulated in Table S2.

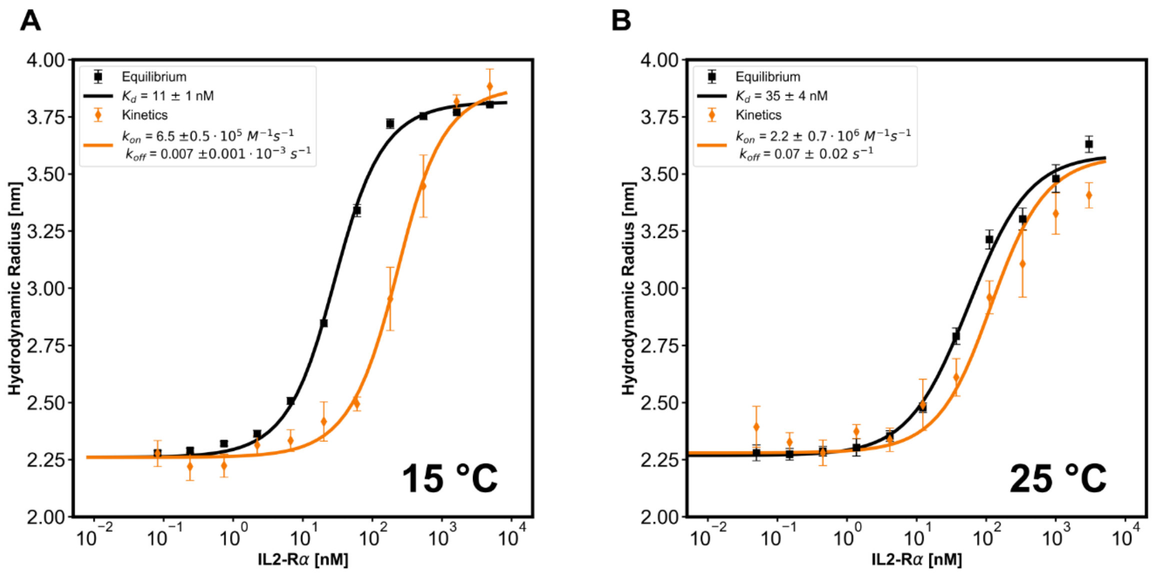


*Figure S5. Interaction of Interleukin-2 with its receptor Interleukin2-Rα in PBS at 15 °C (A) and 25 °C (B).*

*Table S2. Comparison of kinetic parameters obtained from C-Jump in different buffers and temperatures and reported literature value from surface-based method.*

| | $k_{on}$ [$s^{-1}M^{-1}$] | $k_{off}$ [$s^{-1}$] | $K_d$ [nM] |
|---|---|---|---|
| $Serum$ (15 °C) | $(1.9 \pm 0.8) \times 10^6$ | $0.04 \pm 0.01$ | $19 \pm 3$ |
| $PBS$ (15 °C) | $(6.5 \pm 0.5) \times 10^5$ | $0.007 \pm 0.001$ | $11 \pm 1$ |
| $PBS$ (25 °C) | $(2.2 \pm 0.7) \times 10^6$ | $0.07 \pm 0.02$ | $35 \pm 4$ |
| $SPR$ ($Buffer$, 25 °C) | $7.8 \times 10^6$ [5] | 0.24 [5] | $31 \pm 0.7$ [5] |

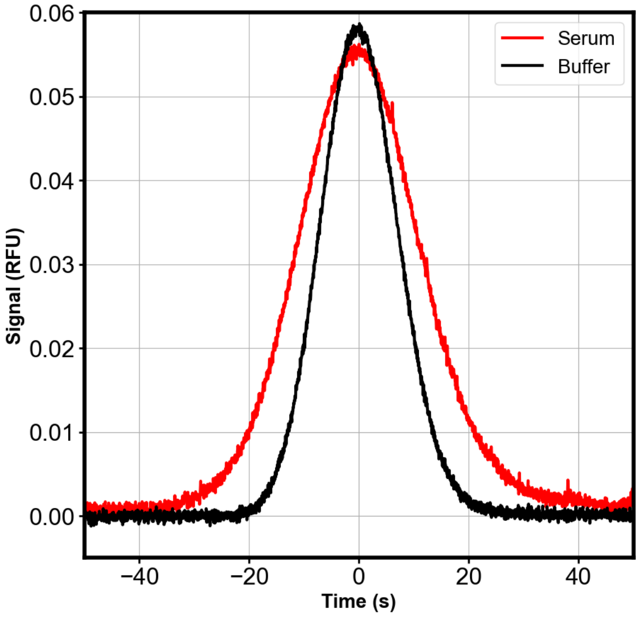


*Figure S6. Recorded fluorescent signals of labeled IL-2 in the absence of analyte, measured in PBS buffer (black) and 90% human serum (red). The serum sample exhibits a broader peak, consistent with increased apparent size due to matrix effects. No secondary peaks or anomalous spikes indicative of nonspecific aggregation or binding are observed.*

### S7 – Setup for ratiometric fluorescence measurement of tryptophan residues

The optical setup follows the same layout as previously described for a red fluorescent dye.[3] Tryptophan residues are excited using an LED at approximately 280nm, and their fluorescence emission is collected between 320 nm and 400 nm. A high-performance dichroic mirror splits the collected light at around 365 nm. The two resulting bands are detected by separate photo detectors, and the fluorescence ratio is calculated by dividing the signal from the longer wavelength range by that of the shorter range. This approach offers high sensitivity, as tryptophan residues are highly responsive to their environment and their emission spectrum can undergo significant changes.[6]

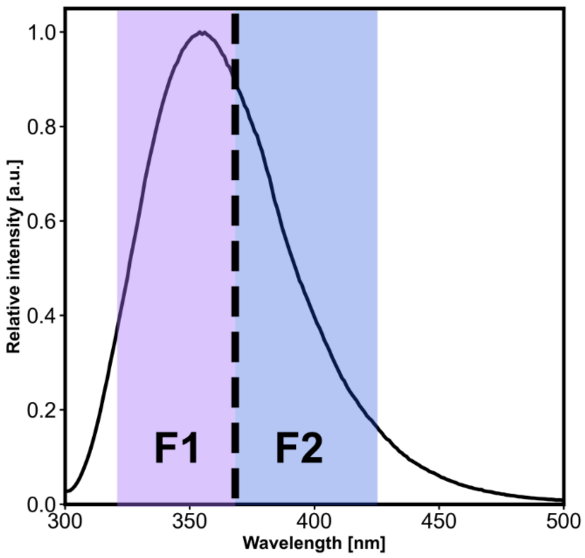


*Figure S7. Emission spectrum of tryptophan.[7] The dashed line indicates where the spectrum is split. F1 and F2 are the resulting spectral bands. The measured ratio is* $\lambda_{2/1} = F_2 F_1^{-1}$.
